# Neutrophil extracellular traps-mediated thrombosis drive pyrrolizidine alkaloid-induced hepatic sinusoidal obstruction syndrome

**DOI:** 10.1101/2025.10.21.683617

**Authors:** Zhang Shuang, Yan Dongming, Cheng Si, Jin Jingyi, Cui Jiamin, Liu Chenghai, Li Yue, Qiu Furong

## Abstract

Hepatic sinusoidal obstruction syndrome (HSOS) caused by pyrrolizidine alkaloid (PA) is a life-threatening disease with limited treatments, yet the initiating vascular event remains undefined. Neutrophil extracellular traps (NETs), as key driver of immunothrombosis in various thromboinflammatory diseases, may represent a critical yet uncharacterized component in the pathogenesis of PA-HSOS. Rats and mice were separately administered monocrotaline (MCT) or senecionine (SEN) via gavage to induce HSOS. Human hepatic sinusoidal endothelial cell (HHSEC) and primary human neutrophil treated with MCT, SEN, or corresponding metabolites were used as in vitro models. In separate experiments, animals received a neutrophil elastase (NE) inhibitor or neutrophil-depleting antibody to evaluate the contribution of NETs. We detected NETs within the sinusoids of necrotic liver lobes, as evidenced by robust immunostaining for NETs markers, accompanied with elevated level of circulating cell-free DNA, and hepatic citrullinated histone H3. Sinusoidal microthrombi containing VWF, CD41, and fibrin(ogen) were also stained positive for neutrophils in PA-HSOS models. Treatment with NE inhibitor or neutrophil depletion markedly attenuated PA-induced liver injury by concomitantly suppressing NETs formation and intrahepatic thrombogenesis. In vitro, conditioned medium from HHSEC exposed to MCT or SEN metabolites potently induced NETs formation. HHSEC injured by PA metabolites released CXCL8. Furthermore, hepatic mRNA and protein levels of CXCL1 and CXCL2 (murine homologous to human CXCL8) were up-regulated in PA-HSOS models, accompanied by a parallel elevation of their high-affinity receptors, CXCR1 and CXCR2. Our study identifies NETs-mediated immunothrombosis as a key driver in HSOS, which provides a promising target for PA-HSOS treatment.

## 1. Introduction

Hepatic sinusoidal obstruction syndrome (HSOS) is a vascular disease, characterized by hepatic sinusoidal congestion and extensive expansion, accompanied with microthrombi formation in sinusoids and post-sinusoidal venules, leading to necrosis of perivenular hepatocytes(DeLeve et al. 2002; Ming 2024). The etiology of HSOS is multifaceted and categorized into three main types: chronic ingestion of pyrrolizidine alkaloids (PAs), use of chemotherapeutic and immunosuppressive agents, and hematopoietic stem cell transplantation (HSCT)(Kumar et al. 2019). PAs and their N-oxides constitute one of the most significant classes of plant-derived phytotoxins. To date, more than 660 distinct compounds have been identified from over 6,000 plant species worldwide, and exposure to PA-containing products have caused large and recurring episodes of acute hepatotoxicity in multiple countries(BL et al. 1999). In Eastern countries, the primary cause of HSOS is ingestion of PA-containing herbals, teas, salads, or PA-contaminated dietary products, including wheat, milk, and honey(Zheng and Zhang 2024). The clinical presentation of PA-HSOS includes ascites, hepatomegaly, jaundice, abdominal distention, and right upper quadrant pain(Barcelos et al. 2021). The mortality rate for PA-HSOS in adults ranges from 16-40%, with liver failure being the leading cause of death(Zheng and Zhang 2024). Currently, therapeutic options have been limited and predominantly focused on supportive or symptomatic management, including liver protection, diuretic therapy, and improvement of microcirculation(Zhuge et al. 2018). While these measurements can attenuate liver injury and promote the recovery of liver function, they seldom produce substantial reversal of pathological and physiological changes in the majority of patients(Zhuge et al. 2019).

Given the high mortality rate of PA-HSOS, extensive efforts have been devoted to characterize the disease. PAs can be metabolized by cytochrome P450 enzymes (CYP450s) into intermediate dehydropyrrolizidine alkaloids (DHPA). DHPAs rapidly interact with cellular macromolecules to form pyrrole-protein adducts (PPAs), which damage the structure of liver sinusoidal endothelial cell (LSEC)(He et al. 2021; Hessel-Pras et al. 2019). The process results in sinusoidal dilation and rupture, with subsequent extravasation of erythrocytes and endothelial cells into the Disse space(Mohty et al. 2015). However, owing to the complexity of its pathogenesis, the mechanisms responsible for the key events remain unclear and greatly hinders the development of effective clinical treatments.

Notably, HSOS can be triggered by diverse etiologies-yet it culminates in highly consistent terminal pathological manifestations, including sinusoidal endothelial cell injury, perisinusoidal fibrosis, obliteration of terminal hepatic venules, and hepatocyte necrosis(Chen and Huo 2010; Mohty et al. 2015). Within this framework, thrombogenesis is regarded as a pivotal event in HSOS pathogenesis: various initial injuries target LSECs, leading to their dysfunction and detachment. This results in exposure of the prothrombotic subendothelial basement membrane and triggers damaged LSECs to secrete abundant pro-coagulant mediators (e.g., VWF) while concomitantly suppressing endothelial thrombomodulin expression(DeLeve et al. 2002). Direct pathological evidence from patient liver tissues further confirms the presence of microthrombi composed of VWF and platelets within the sinusoids(Nishigori et al. 2015). Moreover, the anti-thrombotic agent defibrotide has been established as standard therapy for HSCT-associated HSOS(Corbacioglu et al. 2016). Collectively, these findings underscore the central role of thrombosis in the pathophysiology of HSOS.

Increasing evidences indicate that neutrophil extracellular traps (NETs) contribute to arterial and venous thrombosis as well as inflammation(Martinod and Wagner 2014). NETs, intricate DNA network structures released by neutrophils in response to various damage stimuli, are attached with proteins such as citrullinated histone H3 (CitH3) and myeloperoxidase (MPO). Besides, NETs serve as a scaffold to attract red blood cells, platelets, and procoagulant molecules such as VWF, fibronectin, and fibrinogen(Yang et al. 2020). Recent studies have shown that neutrophils actively participate in the formation of thrombi within blood vessels, a term called immunothrombosis, that can aggravate tissue damage by triggering vessel occlusion and hypoxia(Grover and Mackman 2018). Hepatic NETs release significantly exceeds that in other microvascular beds, as NETs can anchor to sinusoidal walls via VWF and maintain integrity under low-flow conditions(Honda and Kubes 2018). Moreover, Narita’s study reveal that neutrophils participate in the deterioration of hepatic microcirculation in HSOS(Narita et al. 2009). However, it is not clear if NETs contribute to sinusoidal thrombogenesis in PA-HSOS.

To date, monocrotaline (MCT)-induced rat model remains the most widely used and well-characterized one for studying HSOS(Kumar et al. 2019). In parallel, senecionine (SEN), the primary toxin in Gynura segetum (Tusanqi), has been reported to recapitulate the key pathological features of human HSOS in mice(Xiong et al. 2024). In this study, MCT and SEN were separately given to rats and mice by oral gavage to reproduce human HSOS. We hypothesize that NETs constitute the missing link between SEC injury and sinusoidal thrombosis. Furthermore, CXCL8, secreted by damaged liver sinusoidal endothelial cell (LSEC), orchestrates neutrophil chemotaxis and subsequent activation.

## 2. Materials and Methods

### 2.1. Materials

MCT, SEN, and dehydromonocrotaline(DHMC) were obtained from Keluoma Biotech (Chengdu, China), Puruifa Biotech (Chengdu, China), and ChemStrong Scientific (Shenzhen, China), respectively. Anti-granulocyte receptor-1 mAb (Gr-1, clone RB6-8C5, Cat#BE0075) and corresponding isotype controls (Rat IgG2A, Cat#BP0089) were purchased from BioXCell (Lebanon, NH, USA). Sivelestat (SIV) was obtained from Huilun Pharmaceutical (Shanghai, China). ALT and AST activity measurement kits were purchased from Jiancheng Bioengineering Institute (Nanjing, China). PicoGreen kit was provided by UElandy Biotech (Suzhou, China). Optiprep was provided by Stemcell Technologies (Vancouver, CAN). Human hepatic sinusoidal endothelial cell (HHSEC) was obtained from Life-iLab (Shanghai, China). Total RNA extraction kit, reverse-transcription kit, and Real-time PCR kit were bought from Sangon Biotech (Shanghai, China), Accurate Biology (Changsha, China) and Yeasen (Shanghai, China), respectively. Primary antibody for CitH3 (Cat#ab281584) was obtained from Abcam (Cambridge, UK). Antibodies for CXC chemokine receptor 1 (CXCR1, Cat#A16386) and CXCR2 (Cat#A3301) were purchased from Abclonal (Shanghai, China). Antibody for GAPDH (Cat#600004-1) was purchased from Proteintech (Wuhan, China). The providers for ELISA kits were listed as follows: Human: MPO-DNA (Cat#BY-EH112650, Byabscience, Nanjing, China), CXCL1 (Cat#BY-EH112946, Byabscience), CXCL2 (Byabscience, Cat#BY-EH112650), CXCL8 (Youpin, Cat#SYP-H0576, Hangzhou, China). Mouse: CXCL1 (Byabscience, Cat#BY-EM220048), CXCL2 (WeiaoBio, Cat#EM30143XS, Shanghai, China), CXCL5 (Byabscience, Cat#BY-EM227806), CXCL7 (Byabscience, Cat#BY-HS504721). Rat: CXCL1 (LunChangShuo, Cat#ED-30072, Xiamen, China), CXCL2 (LunChangShuo, Cat#ED-31870), CXCL5 (Byabscience, Cat#BY-ER334835), CXCL7 (Byabscience, Cat#BY-HS506051).

### 2.2. Animals

Wild male Sprague-Dawley rats aged 4-6 weeks and male C57BL/6J mice aged 4-6 weeks were purchased from SLAC Laboratory Animal Co., Ltd (Shanghai, China). Rats and mice were bred under specific pathogen-free conditions at Shanghai University of Traditional Chinese Medicine (SUTCM). Animals received standard rodent chow and water ad libitum. All experimental protocols were approved by the Institutional Animal Care and Use Committee of SUTCM (Approval Number: PZSHUTCM2507010001, PZSHUTCM2505200007) and complies with ARRIVE guidelines.

### 2.3. Induction of PA-HSOS in rats and mice

MCT and SEN were firstly dissolved in 0.1 M HCl. The solutions were then neutralized to pH 6.0 with 0.5 M NaOH, and brought to final concentrations of 9 mg·mL^-1^ (MCT) and 6 mg·mL^-1^ (SEN) by adding phosphate buffered saline (PBS). Rats and mice (n=6) were separately gavaged with MCT (90 mg·kg^-1^) or SEN (60 mg·kg^-1^) after overnight fasting to induce PA-HSOS. MCT-induced model in rats is the most common and prevalent HSOS model, and dose of 90 mg/kg was used in most studies(Kumar et al. 2019). It is reported that the median lethal dose (LD50) of senecionine in male C57BL/6J mice is 57.3 mg/kg body weight(Xiong et al. 2012). In our preliminary experiment, C57BL/6J mice received a single SEN gavage at the dose of 50, 60, or 70 mg/kg. Exposing mice to 50 mg/kg SEN did not induce sever liver injury. However, exposing mice to SEN at the dose of 70 mg/kg induced 50% morality within 48 h. In contrast, SEN at a dose of 60 mg/kg caused no death but evoked significant hepatic damage. Consequently, 60 mg/kg was selected as the dose for subsequent mechanistic study of SEN-induced hepatotoxicity. Age-matched rats or mice were gavaged with PBS (pH6.0) and served as vehicle controls. Blood and liver samples were collected 48 h after PAs administration.

### 2.4. Neutrophil depletion in mice

Anti-granulocyte receptor-1 mAb and corresponding isotype control were separately injected i.p. at a dose of 100 μg per mouse 6 hours before and 18 hours after SEN challenge (n=6). Neutrophil depletion efficacy was validated by Wright-Giemsa staining and flow cytometry.

### 2.5. Treatment with pharmacologic inhibitors against neutrophil elastase in rats

Rats (n=6) received an intraperitoneal (i.p.) injection of SIV (a selective inhibitor of neutrophil elastase (NE), 30 mg·kg^-1^) 48 and 24 hours before MCT administration, followed by additional doses immediately after MCT injection (0 h) and at 8-, 24– and 32-hours post-challenge. Control and model animals (n=6) received equivalent volumes of saline at the same time points.

### 2.6. Biochemical assays of blood samples and histology

Both rats and mice were anesthetized and blood samples were collected via the retro-orbital puncture using a heparinized glass capillary tube. Serum activities of ALT and AST were measured using commercially available kits. Paraffin-embedded liver tissues were sectioned (4 μm) for H&E staining to evaluate histologic lesions and coagulation.

### 2.7. Serum cfDNA detection

The assay was performed according to the PicoGreen kit protocol. Briefly, serum samples were diluted 200-fold with PBS and 100 μL was transferred to a black microplate, followed by the addition of 100 μL PicoGreen reagent per well. The plate was incubated at 37℃ in the dark for 5 min. Fluorescence was measured with a microplate reader (Cytation3, BioTek, Winooski, VT, USA) at excitation 480 nm and emission 520 nm. The concentration of each unknown sample was calculated from the standard curve.

### 2.8. Immunofluorescence of NETs and thrombosis

The immunofluorescence of NETs and thrombosis were detected by Tyramide Signal Amplification-based Multiplex Immunofluorescence (TSA-mIF). Briefly, small pieces of liver tissues in OCT were immediately frozen by liquid nitrogen. Frozen sections (8-μm) were prepared using a cryostat microtome (CM3050S, Leica, Wetzlar, Germany) and then stored at –80℃. After fixed with 4% paraformaldehyde, antigen retrieval and endogenous peroxidase blocking, sections were blocked with 5% goat serum. The sections were then incubated with primary antibody for MPO and HRP-labeled secondary antibody. Fluorescent dye TYR-570 was added to produce red fluorescence. Then the MPO antibody was stripped off by microwave, and the sections were incubated with other primary antibodies (Cit-H3, VWF, CD41, Fibrin) plus HRP-labeled secondary antibody, followed by fluorescent dye TYR-520 addition to generate green fluorescence. Nuclei were stained with DAPI. Images were acquired using an inverted fluorescence microscope (IX71, Olympus, Tokyo, Japan; MIX60-FL, Mingmei, Guangzhou, China).

### 2.9. RNA sequencing and bioinformatics analysis

Total RNA extraction, library construction and Illumina RNA-sequencing were outsourced to Beijing Biomarker Technologies Co., Ltd. (Beijing, China), to which liver specimens from both control and model groups were shipped on dry ice. Based on the gene expression matrix, differential expressed analysis was performed in R v4.5.1 with DESeq2 (v1.48.1). P values were adjusted to control the false-discovery rate (FDR) using the Benjamini-Hochberg procedure; |log2 fold-change| ≥ 1 and FDR < 0.05 were considered significant. Volcano plots were generated with ggplot2 (v4.0.0). Functional enrichment was assessed against the Kyoto Encyclopedia of Genes and Genomes using clusterProfiler (v4.16.0). Enrichment P values were calculated by hypergeometric test, adjusted for multiple testing (Benjamini–Hochberg); pathways with FDR < 0.05 were deemed significant. For visualization of enriched pathways, z-score scaled mean expression values of differential expression genes within pathway of interest were extracted and plotted as heat maps using pheatmap (v1.0.13).

### 2.10. Culture and treatment of HHSEC

HHSEC was seeded at a density of 2×10^5^ cells/mL in T25 cell culture flasks and cultured overnight in a humidified incubator at 37℃ with 5% CO₂. Then MCT or DHMC (both at a final concentration of 250 μM) was added to the corresponding flask. Following a 24-hour treatment, supernatant was harvested and used as conditioned media in subsequent assays. The concentrations of MCT and DHMC were selected on the basis of the study by Yang et al., in which 300 μM DHMC evoked profound GSH depletion and time-dependent (6–48 h) cytotoxicity in HHSEC(Yang et al. 2016).

### 2.11. Isolation of primary human neutrophils and platelets

Primary human neutrophils (PMNs) were isolated using polysucrose sedimentation, Optiprep density gradient centrifugation, and erythrocyte hypotonic lysis following standard procedures as provided by manufacture. The number and purity of neutrophils in each sample was detected using an automated hematology analyzer (XT-2000i, Sysmex, Kobe, Japan). Human platelets were isolated from whole blood by centrifugation. Briefly, anticoagulated whole blood was centrifuged at 800 rpm to prepare platelet-rich Serum, followed by a second centrifugation at 3000 rpm to pellet the platelets. The studies were approved by the Research Ethics Committee of Shuguang Hospital affiliated to Shanghai University of Traditional Chinese Medicine (Approval Number: 2025-1856-196-02).

### 2.12. Induction of NETs formation in neutrophils culture alone

Freshly isolated human primary neutrophils were seeded into a 48-well plate at a density of 5×10^5^ cells/mL. After allowing for adherence for approximate 15 minutes, the medium was replaced with MCT, DHMC, supernatant from MCT-treated HHSEC, supernatant from DHMC-treated HHSEC, or Phorbol 12-myristate 13-acetate (PMA) as positive control. The final concentrations of MCT, DHMC, and PMA were 250 µM, 250 µM, and 100 ng/mL, respectively. After a further 4-hour incubation, the supernatant was collected. The concentration of cell-free DNA (cfDNA) and MPO-DNA in the supernatant was determined using PicoGreen reagent and an ELISA kit, respectively. To distinguish cell-free NETs that were released away from neutrophils and anchored NETs that were anchored to neutrophils, both supernatant and neutrophils anchored to the plate were collected for SytoxGreen staining. The supernatant was transferred to a new 48-well plate, and then SytoxGreen and Hoechst 33342 (at final concentrations of 5 µM and 10 µg/mL, respectively) were added. After incubation for 30 minutes, cell-free NETs were visualized using fluorescence microscopy. In parallel, SytoxGreen and Hoechst at the same concentrations were added to anchored neutrophils and immunofluorescence was detected.

### 2.13. Induction of NETs formation in neutrophils co-cultured with platelets

Freshly isolated neutrophils and platelets were co-cultured in a 48-well plate at densities of 5×10^5^ cells/mL and 1×10^7^ cells/mL, respectively. After 15-min attachment, the medium was replaced with MCT, DHMC, or conditioned medium harvested from MCT– or DHMC-treated cells. The final concentrations of MCT and DHMC were both 250 µM. After a further 4-hour incubation, the supernatant was collected. The concentration of cfDNA and MPO-DNA, as well as the level of cell-free NETs and anchored NETs, were detected as the aforementioned descriptions.

### 2.14. Metabolic study of SEN in human liver microsomes

The human liver microsome (HLM) reaction system was composed as follows: HLMs (0.5 mg/mL), MgCl₂ (3.2 mM), glutathione (GSH, 10 mM), SEN (250 μM), and PBS. After pre-incubation at 37℃ for 5 minutes, the reaction was initiated by adding nicotinamide adenine sinucleotide phosphate (NADPH, 1 mM) with a total volume of 60 μL. After a 2-hour incubation, 60 μL of ice-cold acetonitrile was added to terminate the reaction. The mixture was vortexed for 5 minutes and then centrifuged at 12,000 rpm for 10 minutes. A 100 μL aliquot of the supernatant was collected and mixed with 200 μL of acetonitrile containing internal standard S-hexylglutathione (SHG). The mixture was analyzed by LC-MS/MS in multiple reaction monitoring (MRM) mode to detect the GSH conjugate of (±)-6,7-dihydro-7-hydroxy-1 –hydroxymethyl-5H-pyrrolizidine (DHP), which was further confirmed in neutral loss (NL) mode. The detailed LC-MS/MS method is provided in the supplementary data. To optimize the concentration of SEN for subsequent studies, enzyme kinetic experiment was conducted. The reaction system was composed as previously described, with SEN concentrations ranging from 8.2 to 2000 µM and reaction time of 20 minutes. The formation rate of DHP-GSH was determined using MRM mode. The Michaelis-Menten equation was used to fit the data, with the formation rate of DHP-GSH plotted on the y-axis and the concentration of SEN on the x-axis, to calculate the kinetic parameters K_m_ and V_max_.

### 2.15. Effects of supernatant from SEN metabolites-treated HHSEC on NETs formation

HHSEC was seeded at a density of 2×10^5^ cells/mL in a 48-well plate and cultured overnight in a humidified incubator at 37℃ with 5% CO₂. Dehydroheliotrine (reactive metabolite of SEN) was generated using HLM reaction system. The reaction system was composed as follows: HLMs (0.5 mg/mL), MgCl₂ (3.2 mM), NADPH (1 mM), SEN (250 μM) in complete HHSEC medium with a total volume of 1 mL. The reaction system without SEN was also prepared. Both systems were incubated at 37℃ for 2 hours. The reaction system with or without SEN was separately added to HHSEC. After a further 12-hour incubation, the supernatant was collected. Freshly isolated neutrophils were seeded at a density of 5×10^5^ cells/mL in a 48-well plate. After allowing for adherence, the culture medium was removed and replaced with 0.2 mL of the three types of HHSEC supernatant collected above. Neutrophils were cultured for 4 hours, and then the supernatant was collected. The concentration of MPO-DNA in the supernatant was measured as described in the kit instruction.

### 2.16. RNA isolation and Real-Time Quantitative PCR

Total RNA in tissues was extracted with Trizol and purified with spin column following the manufacturer’s instructions. Purified total RNA was reverse-transcribed into complementary DNA using the PrimeScript RT reagent kit. Real-time PCR was performed using qPCR SYBR Green Master Mix with a primer set (sequences are listed in Supplementary Table. 1) on a real-time PCR detection system (Viia 7, Applied Biosystems, Foster City, CA, USA).

### 2.17. Immunoblotting analysis

Liver proteins were extracted with RIPA buffer supplemented with 1 mM PMSF, resolved by SDS-PAGE, and transferred to PVDF membranes. The membranes were then probed with primary antibodies against CitH3, CXCR1, CXCR2 or GAPDH, followed by incubation with corresponding secondary antibodies conjugated with horseradish peroxidase. Protein bands were visualized with ECL substrate (Genebrick, Shanghai, China) in a gel imaging system (Biorad, Hercules, CA, USA). ImageJ software (NIH, Bethesda, MD, USA) was used to analyze the optical density of protein bands and normalize them to GAPDH.

### 2.18. ELISA assay

Chemokine levels in supernatant from HHSEC and in liver homogenates from rodents were determined by enzyme-linked immunosorbent assay (ELISA). For HHSEC, conditioned medium was collected and centrifuged at 12,000 rpm for 5 min. Then the resulting supernatant was stored at –80 °C until analysis. For liver tissues, 100 mg of mouse or rat livers was homogenized in 900 µL ice-cold PBS containing 1× protease-inhibitor cocktail. The homogenate was centrifuged at 10,000 rpm for 10 minutes at 4 °C, and then the supernatant was collected. All assays were performed according to the manufacturers’ protocols. Protein concentrations in liver homogenates were measured in parallel using bicinchoninic acid assay (BCA) assay. Chemokine concentrations were normalized to milligrams of total protein and expressed as pg·mg⁻¹ protein.

### 2.19. Data analysis

To confirm the presence of hepatic sinusoidal thrombosis, we developed a fluorescence-based assay that measures the spatial distances between MPO and CitH3, CD41, VWF, and fibrin(ogen), thereby clarifying neutrophil involvement in thrombus formation. All fluorescence images used for comparison were re-scaled to identical pixel dimensions and then pre-processed by ImageJ to eliminate background fluorescence interference. Total fluorescence intensities of CitH3, CD41, VWF, and fibrin(ogen) were measured using the Measure Image Intensity module in CellProfiler software (Broad Institute, Cambridge, MA, USA). Fluorescent puncta corresponding to MPO, CitH3, CD41, VWF, and Fibrin(ogen) were identified using the Indentify Primary Objects module in CellProfiler, and their centroid coordinates were recorded. The centroid coordinates of all MPO puncta constituted the reference set, whereas the centroid coordinates of the remaining puncta (CitH3, CD41, VWF and fibrin(ogen)) served as query sets. For every punctum, the Euclidean distance to its nearest MPO punctum was calculated with the RANN package in R.

Quantitative data are presented as mean±SD. Statistical analyses were performed using the GraphPad Prism 8 (GraphPad, San Diego, CA, USA). Statistical differences between two groups were analyzed by an unpaired 2-tailed Student’s t test or t-test with Welch’s correction for variables with or without the same variability of difference, respectively. Differences among multiple groups were assessed by one-way analysis of variance (one-way ANOVA) with or without Welch’s correction based on whether the Brown-Forsythe test P-value is less than 0.05 or not. In all statistical comparisons, *p*<0.05 was used to indicate a significant difference.

## 3. Results

### 3.1. NETs form in liver of PA-HSOS models

In this study, two well-characterized murine models of HSOS were used-rats administered MCT and mice receiving SEN (Fig. 1A, H). Compared with normal ones, model animals exhibited deep-red, congested livers with increased firmness (Fig. 1B, L), and serum ALT and AST levels were markedly elevated (Fig. 1C, J). H&E staining revealed that livers from control rats and mice displayed intact central veins and portal areas. Each hepatic lobule has a central vein and radiated strands separated by sinusoids lined with endothelial cells. Contrarily, livers from model animals showed dilated and distorted sinusoids, sinusoidal spaces occluded by erythrocytes, massive neutrophils infiltration within the sinusoids, and hepatocyte coagulative necrosis (Fig. 1D, K).

**Fig. 1.**
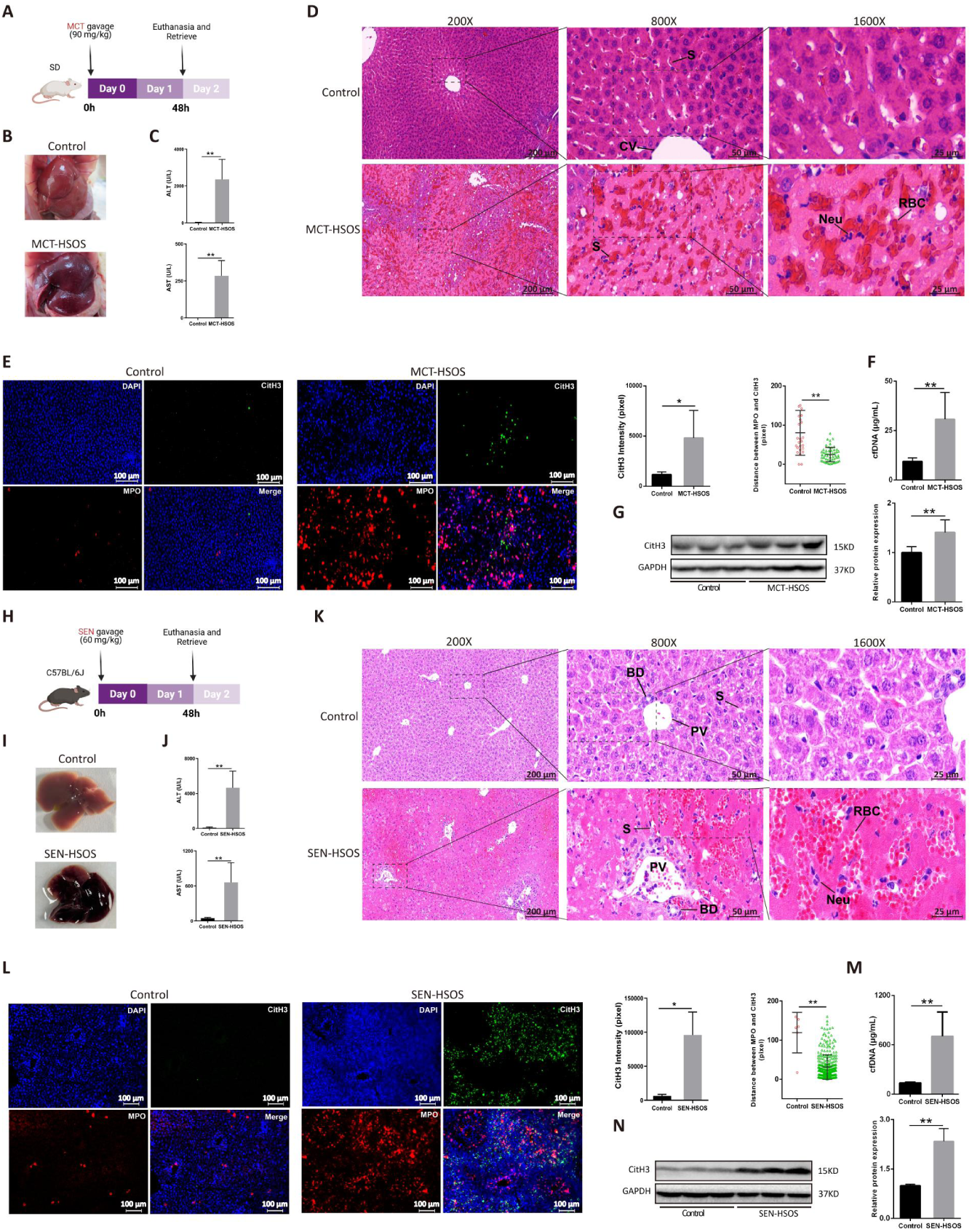
NETs are present in the PA-HSOS models. (A-I) MCT-HSOS rats: A. Experimental scheme of MCT-HSOS in rats. B. Gross liver morphology. C. Serum ALT and AST activities. D. Representative H&E-stained liver sections. E. Multiplex IF images of liver tissues stained for MPO and CitH3, and corresponding CitH3 quantification and proximity analysis. F. Serum cfDNA level. G. Hepatic CitH3 protein expression. (H-N) SEN-HSOS mice: H. Experimental scheme of SEN-HSOS in mice. I. Gross liver morphology. J. Serum ALT and AST activities. K. Representative H&E-stained liver sections. L. Multiplex IF images of liver tissues stained for MPO and CitH3, with CitH3 quantification and proximity analysis. M. Serum cfDNA level. N. Hepatic CitH3 protein expression. CV: central vein; S: sinusoid; PV: portal vein; BD: bile duct; Neu: Neutrophil; RBC: red blood cell. Data are expressed as mean±SD, \*\**p*<0.01, \**p*<0.05 versus control groups.

To determine whether infiltrating neutrophils form NETs, immunofluorescence was employed to detect the co-localization of MPO, CitH3, and DAPI. Our results revealed sparse neutrophils in hepatic sinusoids of control mice and rats, whereas both MPO and CitH3 signals were markedly intensified in HSOS models, with CitH3 foci frequently adjacent to or overlapping MPO-positive neutrophils-an observation consistent with robust NETs formation (Fig. 1E, L). Circulating cfDNA levels and hepatic CitH3 protein expression corroborated these findings: serum cfDNA was 3.3-fold (MCT-HSOS) and 5.2-fold (SEN-HSOS) higher versus controls (Fig. 1F, M), and hepatic CitH3 protein rose to 1.4-fold and 2.3-fold of control group, respectively (Fig. 1G, N). Collectively, these results suggest NETs formation by infiltrating neutrophils in both PA-HSOS models.

### 3.2. Immunothrombosis containing neutrophils exist in PA-HSOS

To delineate the existence of immunothrombosis in HSOS, we employed mIF to map the cellular compositions of hepatic microthrombi. Sinusoids were considered to contain immunothrombosis if there was a occlusion of the sinusoid that stained positive for a network of neutrophils(MPO), VWF, fibrin(ogen), or platelets(CD41)(Nicolai et al. 2020). VWF was predominantly present in vascular endothelial cells in control group. Contrarily, VWF expression was markedly elevated and distributed diffusely along the sinusoidal lining in PA-HSOS models. The results also revealed numerous intravascular neutrophils in closer association with VWF (68.9 vs 43.4 pixels in rats; 188.3 vs 53.8 pixels in mice), indicating neutrophils adhesion to VWF (Fig. 2A). We also detected the presence of neutrophils in fibrin clot, as corroborated by strongly increased fibrin(ogen) signal and closer distance between MPO and fibrin(ogen) in sinusoids (30.9 vs 26.6 pixels in rats; 22.5 vs 20.0 pixels in mice) (Fig. 2B). Notably, both MCT– and SEN-induced HSOS models showed markedly closer MPO-CD41 signals (46.6 vs 27.2 pixels in rats; 54.5 vs 34.3 pixels in mice), indicating neutrophil-platelet aggregates formation (Fig. 2C).

**Fig. 2.**
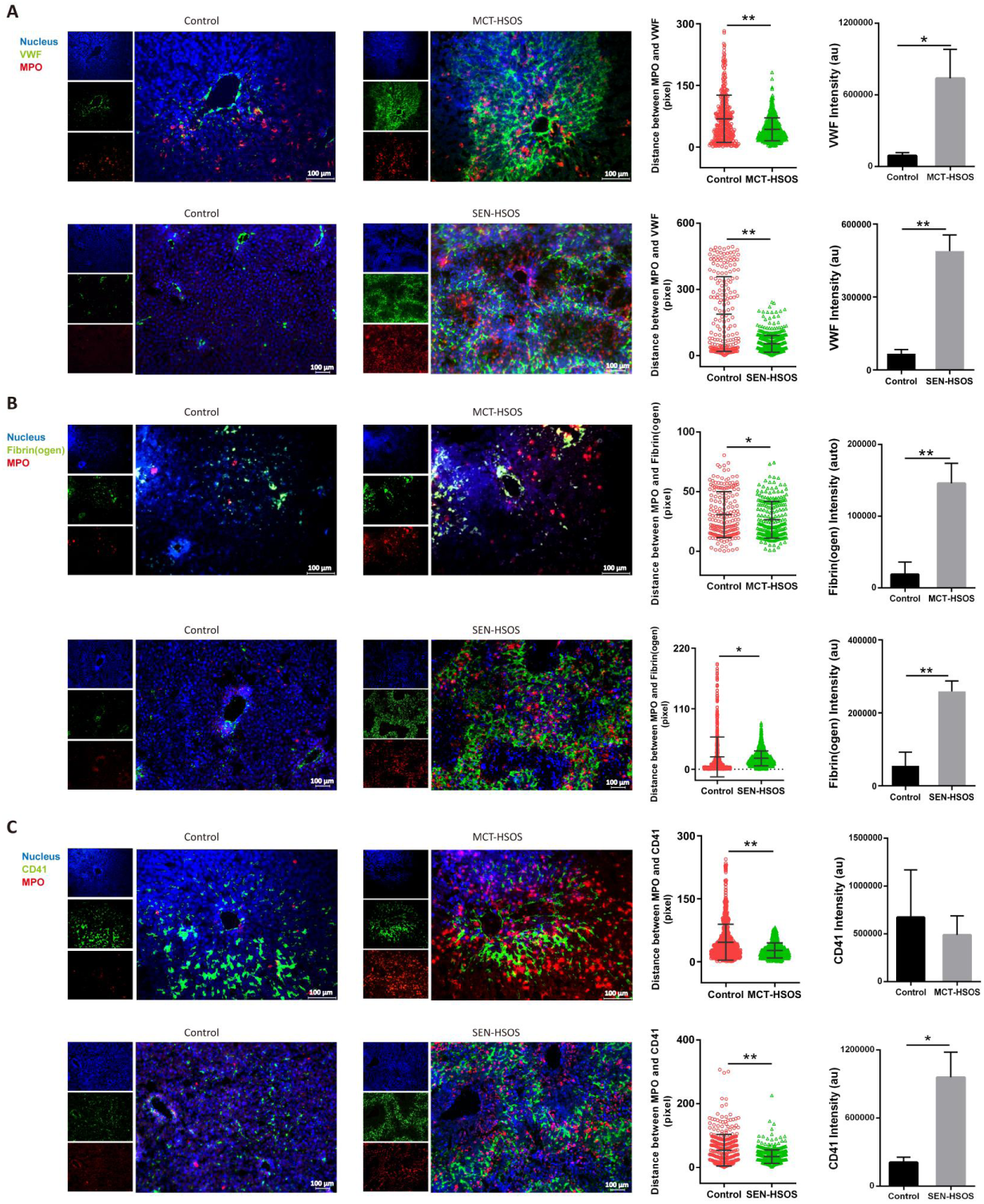
Immunothrombi form in PA-HSOS models. Representative multiplex IF images of hepatic microthrombi showing: A. MPO with VWF, including proximity analysis and VWF quantification. B. MPO with fibrin(ogen), including proximity analysis and fibrin(ogen) quantification. C. MPO with platelet glycoprotein CD41, including proximity analysis and CD41 quantification. Data are expressed as mean±SD, \*\**p*<0.01, \**p*<0.05 versus control groups.

### 3.3. Inhibit pro-thrombotic NETs attenuates liver injury induced by PA

Next, the anti-mouse Gr-1 mAb was used to deplete neutrophils in mice, and depletion efficacy was validated by Wright-Giemsa staining and flow cytometry (Fig. 3A, Supplementary Fig. 1). Depletion of neutrophils substantially ameliorated hepatic sinusoidal congestion, sinusoidal architecture disruption, and hepatocellular coagulative necrosis (Fig. 3B). Concomitantly, Serum ALT and AST concentrations were reduced to 43.6 and 51.7% of the model group, respectively (Fig. 3C). Using the above-mentioned mIF assay, we confirmed that NETs was reduced following neutrophil ablation (Fig. 3D). Furthermore, fluorescence intensities of VWF and fibrin(ogen) were diminished, correlated with longer distance between neutrophil and platelet, or fibrin(ogen), indicating that neutrophil depletion attenuates HSOS-induced thrombosis (Fig. 3E, F, G).

**Fig. 3.**
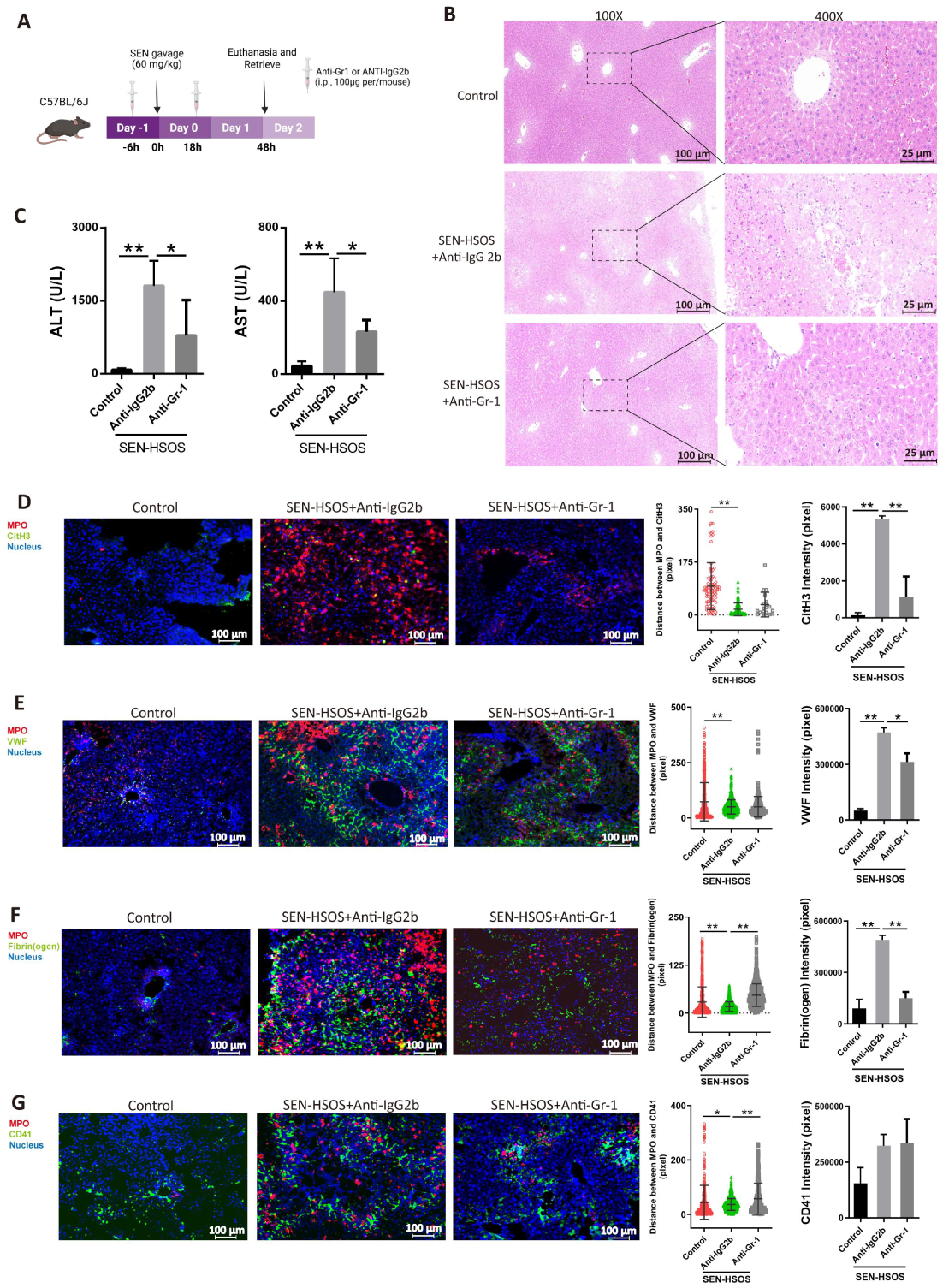
Liver injury is ameliorated in SEN-HSOS model with neutrophil depletion. A. Experimental scheme of the experiment. B. Representative images of H&E-stained liver sections. C. Serum ALT and AST activities. D. Immunofluorescence of CitH3 and MPO in liver sections, and corresponding proximity analysis and CitH3 quantification. E. Immunofluorescence of VWF and MPO in liver sections, and corresponding proximity analysis and VWF quantification. F. Immunofluorescence of fibrin(ogen) and MPO in liver sections, and corresponding proximity analysis and fibrin(ogen) quantification. G. Immunofluorescence of CD41 and MPO in liver sections, and corresponding proximity analysis and CD41 quantification. Data are expressed as mean±SD, \*\**p*<0.01,\**p*<0.05.

To verify our findings derived from neutrophil depletion, we next employed pharmacologic NE inhibition on MCT-induced HSOS (Fig. 4A). Sivelestat (SIV), a selective NE antagonist, effectively suppresses NETs formation and has been clinically applied in conditions such as acute lung injury(Ren et al. 2024). Rats treated with SIV before and after MCT administration exhibited markedly attenuated liver injury, with serum ALT and AST levels reduced to 38.0 and 22.5% of model group, respectively (Fig. 4B). SIV treatment largely reversed hepatic pathological patterns, preserving vascular architecture, limiting erythrocytes extravasation, and ameliorating necrosis (Fig. 4C). Moreover, liver from SIV-treated mice demonstrated less NETs formation compared with those in model group (Fig. 4D). SIV treatment significantly reduced VWF expression and increased the intercellular distance between neutrophils and platelets (from 25.91 to 36.80 pixels), indicating reduced VWF fiber and neutrophil-platelet aggregates formation (Fig. 4E, F). These results verify the critical role of NETs in driving thrombosis in MCT-induced HSOS.

**Fig. 4.**
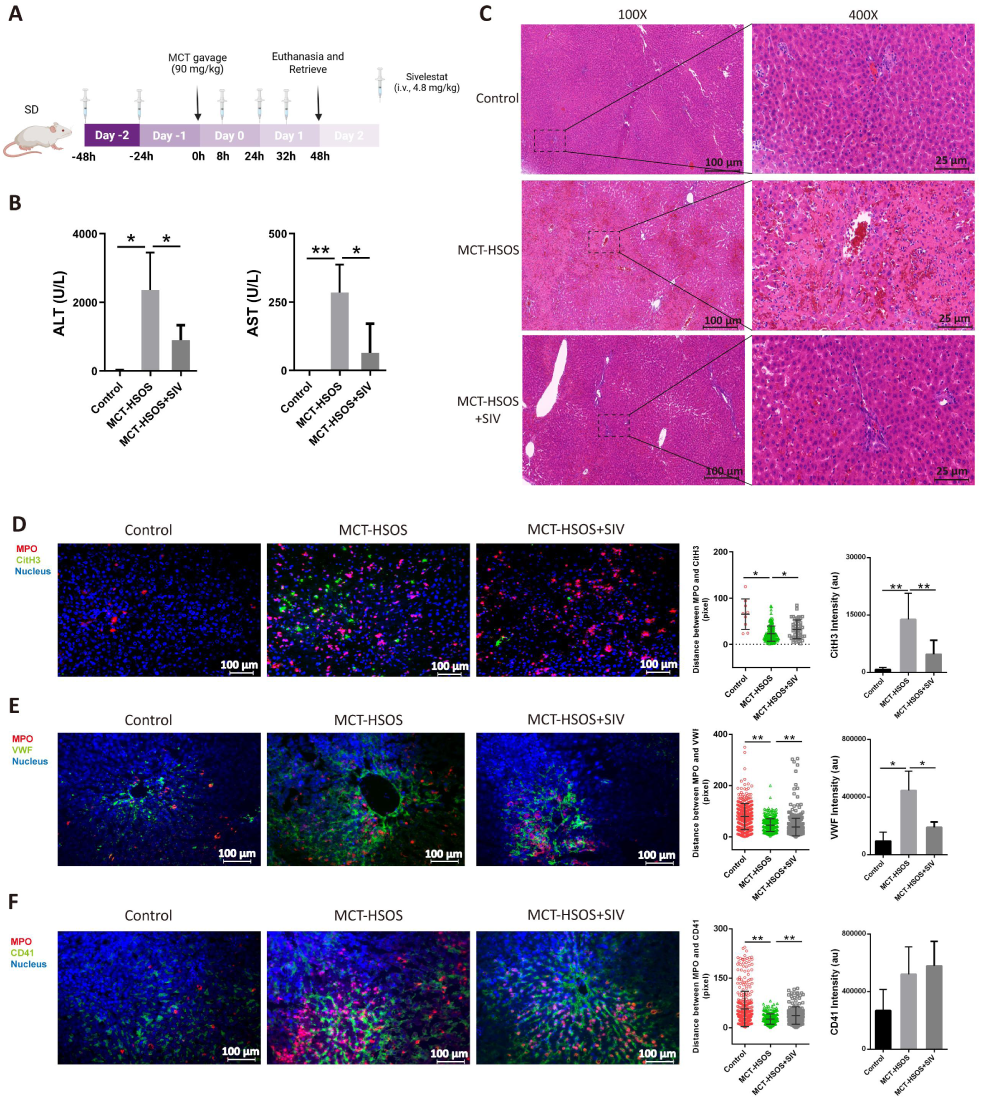
Sivelestat protects against MCT-induced liver injury. A. Experimental scheme of the experiment. B. Serum ALT and AST activities. C. Representative image of H&E-stained liver sections. D. Immunofluorescence of CitH3 and MPO in liver sections, with proximity analysis and CitH3 quantification. E. Immunofluorescence of VWF and MPO in liver sections, with proximity analysis and VWF quantification. F. Immunofluorescence of CD41 and MPO in liver sections, with proximity analysis and CD41 quantification. Data are expressed as mean±SD, \*\**p*<0.01, \**p*<0.05.

### 3.4. Conditioned medium from PA metabolites-treated HHSEC stimulates neutrophil activation

We therefore accessed NETs formation in four *in vitro* conditions. Firstly, neutrophils were exposed to MCT or its metabolite DHMC, with PMA as a positive control. Neither MCT nor DHMC increased cfDNA levels, and the concentration of MPO-DNA did not reach the threshold to be detected (<1.25 ng·mL^-1^). SytoxGreen fluorescence in supernatant or adherent cells did not differ from baseline. PMA-treated neutrophils exhibited no cell-free NETs yet displayed a robust SytoxGreen signal in adherent cells (Fig. 5A). Secondly, Neutrophils and platelets were co-incubated with MCT or DHMC. Again, no cell-free NETs were detected. MPO-DNA remained at baseline and below quantifiable levels, and SytoxGreen staining was negligible (Fig. 5B).

**Fig. 5.**
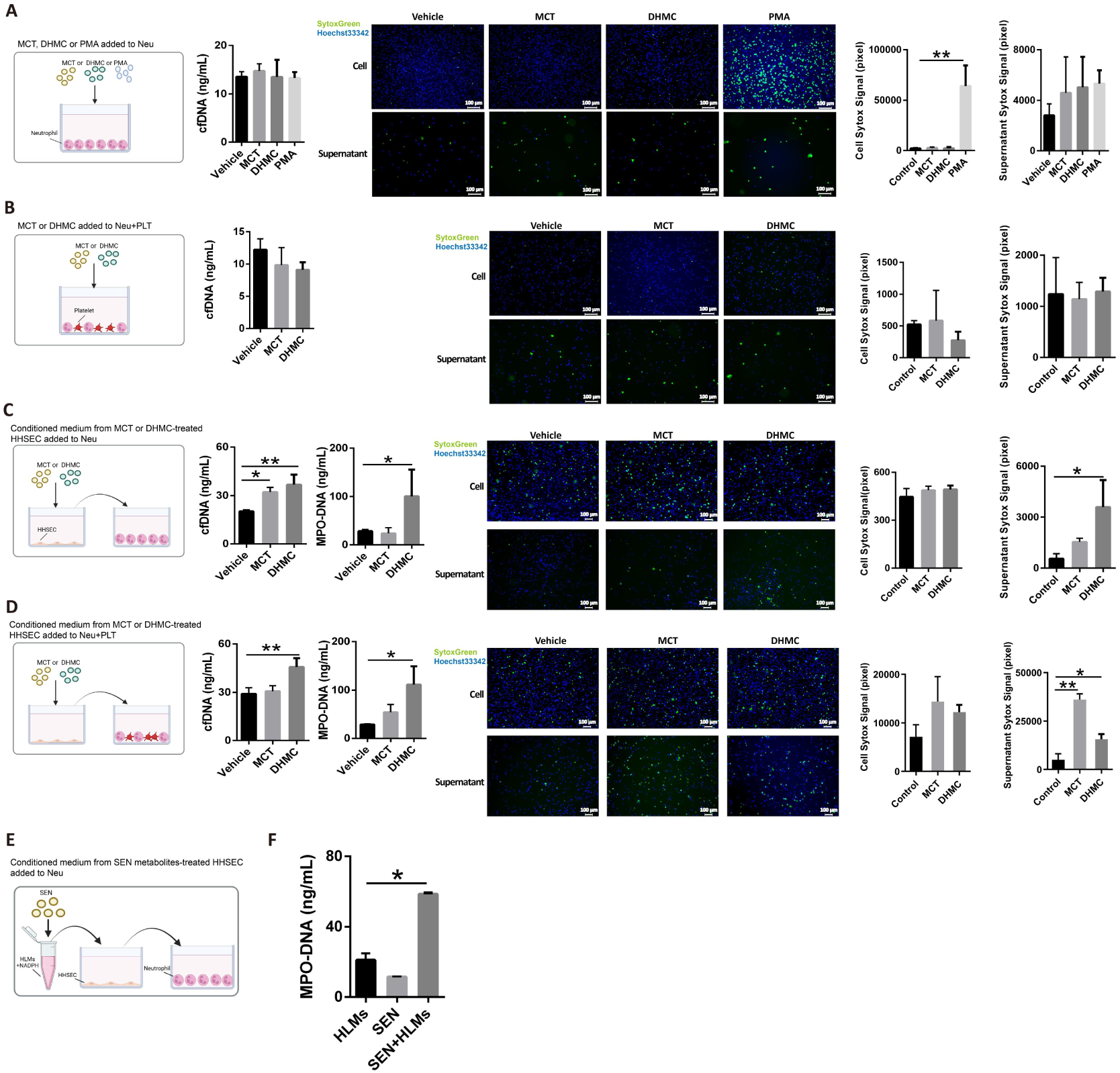
PA metabolites-treated HHSEC supernatant induces NETs formation. A. Neutrophils directly treated by MCT, DHMC, or PMA. B. Neutrophil-platelet co-culture treated by MCT or DHMC. C. Conditioned medium from MCT– or DHMC-treated HHSEC transferred to neutrophils. D. Conditioned medium from MCT– or DHMC-treated HHSEC transferred to neutrophil-platelet co-culture. In all conditions, supernatant cfDNA levels, as well as representative images and intensity of SytoxGreen in cells and supernatant are shown. E. Conditioned medium from HHSEC stimulated with SEN in co-culture with HLMs was transferred to neutrophils. The MPO-DNA levels in the supernatant of conditions A and B are below the lower limit of quantification (LLOQ), and are therefore not shown. Data are expressed as mean±SD, \*\**p*<0.01,\**p*<0.05 versus vehicle groups.

Thirdly, conditioned medium from MCT– or DHMC-treated HHSEC was transferred to neutrophils. DHMC conditioned medium significantly elevated neutrophil supernatant cfDNA and MPO-DNA levels, as well as SytoxGreen intensity. Neutrophils exposed to DHMC conditioned medium exhibited a different profile from that in PMA-treated ones: a strong extracellular fluorescence in the supernatant yet persistent membrane impermeability to SytoxGreen, confirming that the cells remained intact while releasing NETs in a vital manner (Fig. 5C). Fourthly, in neutrophil-platelet co-culture treated with the same conditioned medium, only DHMC conditioned medium elicited cell-free NETs, with cfDNA and MPO-DNA levels comparable to those observed in the absence of platelets (Fig. 5D). Overall, the results indicate that DHMC-injured HHSEC conditioned medium selectively raised cell-free NETs.

Due to a lack of commercially available dehydroheliotrine, SEN was co-incubated with HLMs to generate its metabolites. Then the mixture was applied to HHSEC, and the conditioned medium was transferred to neutrophils (Fig. 5E). LC-MS/MS in MRM and NL modes confirmed formation of the DHP-GSH conjugate. Enzyme kinetic analysis showed K_m_=204.7 μM and V_max_=0.010 min⁻¹ for DHP-GSH generation (Supplementary Fig. 2). MPO-DNA level was markedly elevated only when neutrophils were exposed to medium from SEN metabolites-stimulated HHSEC, indicating induction of NETs (Fig. 5E).

### 3.5. CXCL8-CXCR1/2 axis drives NETs formation in PA-HSOS

RNA-seq profiling of liver tissue detected 15,395 expressed genes. Compared with controls, PA-HSOS livers exhibited 2,090 up-regulated and 2,487 down-regulated transcripts (Fig. 6A). These differentially expressed genes (DEGs) were significantly enriched in 572 KEGG pathways; among the top 30, “cytokine-cytokine receptor interaction” (mmu04060) displayed the strongest association with the robust neutrophil infiltration and NETs observed histologically (Fig. 6B). Heat-map visualization of DEGs within this pathway revealed marked induction of neutrophil chemoattractants CXCL1/2/3/5/7) and their cognate receptor CXCR2 (Fig. 6C), indicating chemokine receptor signaling as a potential driver of NETs during PA-HSOS.

**Fig. 6.**
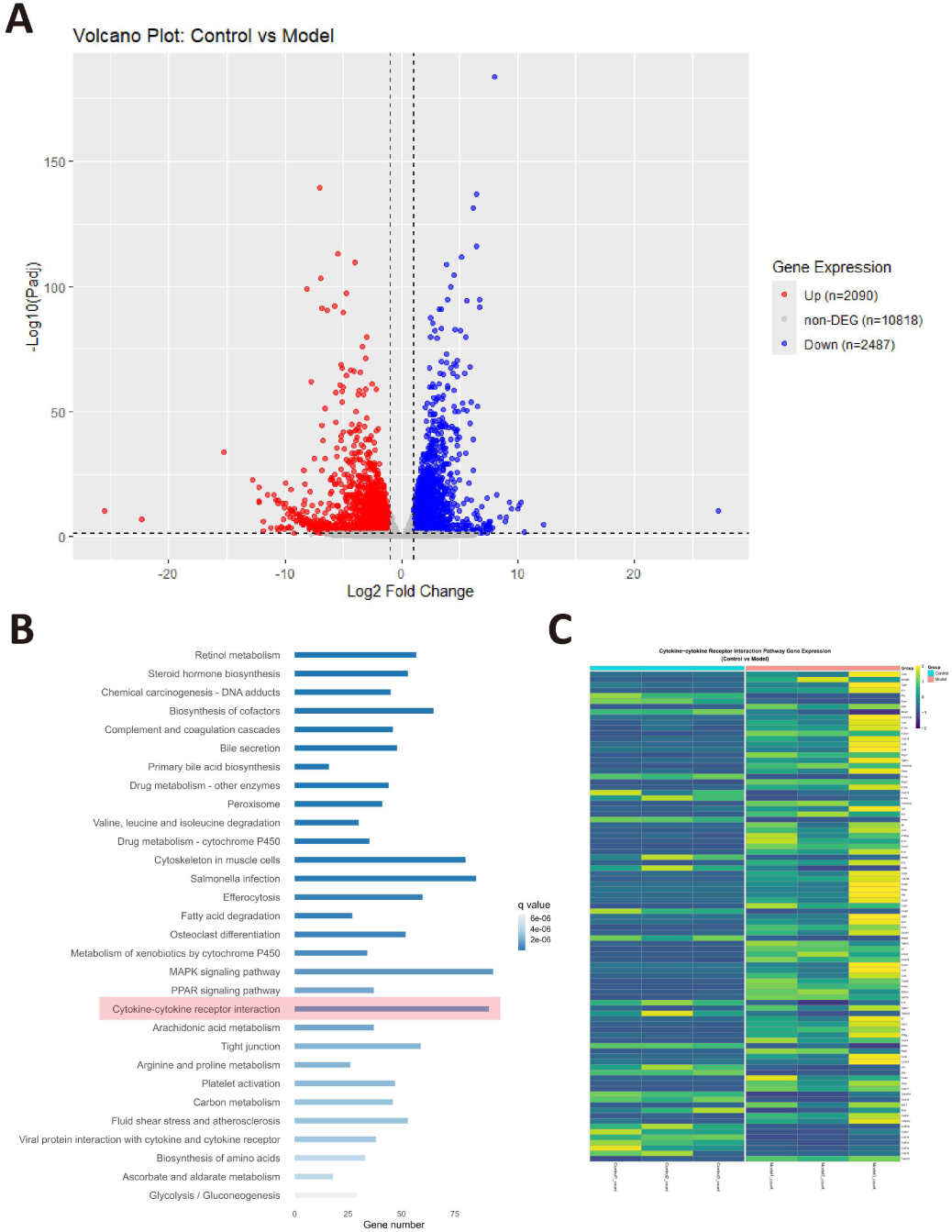
Comparative transcriptomic profiling of control versus SEN-HSOS model livers. A. Volcano plot of differentially expressed genes (DEGs) between control and model groups. B. Top 30 KEGG pathways enriched among DEGs ranked by q value; C. Heatmap showing z-score scaled expression of DEGs involved in cytokine-cytokine receptor interaction pathway.

Next, we measure mRNA levels of the functional homologues (CXCL1/2/3/5/7) and verified the corresponding protein expression in liver. In SEN-HSOS mice, hepatic mRNA of CXCL1, CXCL2, CXCL5 and CXCL7 was markedly up-regulated (Fig. 7A). Concomitantly, CXCL1 and CXCL2 proteins increased to 1.86– and 1.33-fold of control ones, respectively (Fig. 7B). Immunoblotting analyses further revealed significant up-regulation of phosphorylated CXCR1 (∼70 kDa) and phosphorylated CXCR2 (∼46 kDa) in the same model, while the fast-migrating phosphorylated CXCR1 (∼55 kDa) was down-regulated (Fig. 7C). Similarly, MCT-HSOS rats showed 1.31– and 3.68-fold rises in hepatic CXCL1 and CXCL2 levels. In parallel, the expression of CXCL5 and CXCL7 rose to 3.19– and 2.38-fold of the control ones (Fig. 7D). CXCL5 and CXCL7 are both released from platelets and bound by CXCR2(Gollomp et al. 2018), indicating a role for platelet-derived chemokines in directing neutrophil migration in MCT-HSOS. Also, the expression of phosphorylated CXCR1 isoforms (∼70 kDa and ∼55 kDa) as well as phosphorylated CXCR2 (∼46 kDa) was up-regulated in MCT-HSOS models, compared with control ones (Fig.7E). Further, we observed DHMC-treated HHSEC displayed a selective induction of CXCL8 secretion, with no discernible effect on CXCL1 or CXCL2 release (Fig. 7F). Collectively, the results indicate that PA metabolites could induce HHSEC to release CXCL8(CXCL1/2 in rodents), which attracts neutrophils by binding CXCR1/2.

**Fig. 7.**
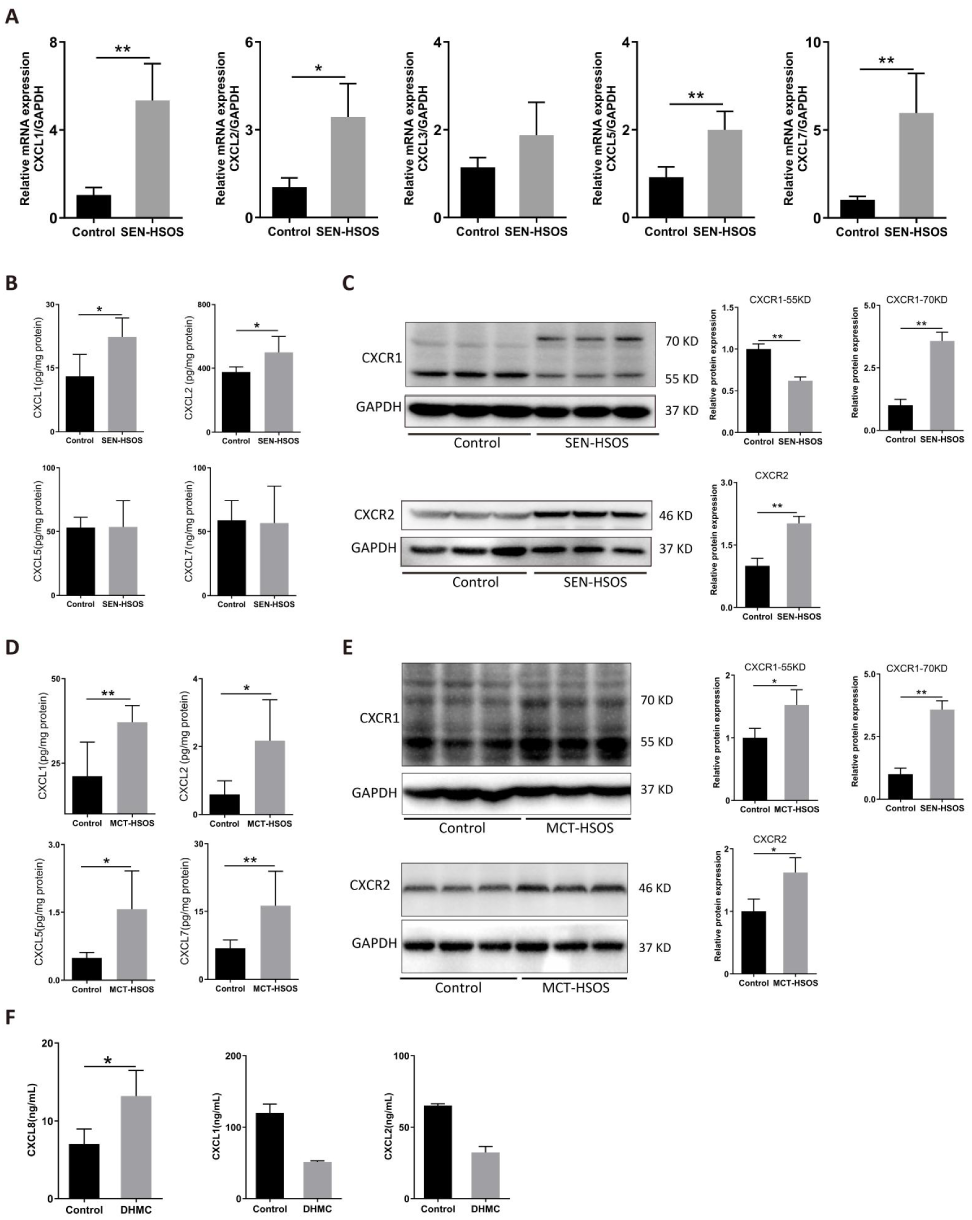
CXCL1/2-CXCR1/2 axis is involved in NETs formation in PA-HSOS. A. Hepatic mRNA level of neutrophil chemokines in SEN-induced HSOS mice. B. Hepatic protein level of neutrophil chemokines in SEN-HSOS mice. C. Hepatic expression of CXCR1 and CXCR2 receptors in SEN-HSOS mice. D. Hepatic protein level of neutrophil chemokines in MCT-induced HSOS rats. E. Hepatic protein level of CXCR1 and CXCR2 in MCT-induced HSOS rats. F. Supernatant chemokine levels in DHMC-treated HHSEC. Data are expressed as mean ± SD, \*\**p*<0.01,\**p*<0.05.

## 4. Discussion

Our study demonstrates NETs formation in both MCT– and SEN-induced HSOS models. By depleting neutrophils and inhibiting NE to reduce NETs, we confirm that NETs are critical contributors to microvascular thrombosis and liver injury. Furthermore, damaged hepatic sinusoidal endothelial cells release CXCL8, which drives neutrophil chemotaxis and subsequent NETs formation.

In the present study, two distinct murine models of HSOS were established, characterized by sinusoidal destruction, erythrocytes extravasation and hepatocyte coagulative necrosis. We observed a marked increase in both MPO and CitH3 signals within the liver in both HSOS models; their pronounced co-localization indicates robust NETs formation. This was corroborated by elevated serum cell-free DNA and increased hepatic CitH3 protein levels, providing independent confirmation of intrahepatic NETs release. NETs serve as a frontline defense by immobilizing and eliminating pathogens. However, excessive NETs formation drives immunothrombosis, disrupting microcirculation and precipitating organ injury(Marcos-Jubilar et al. 2023). We find co-localization of neutrophils and red blood cells in sinusoidal thrombi in PA-HSOS. The immunofluorescence also shows numerous intravascular neutrophils in closer association with fibrin, VWF, and CD41, indicating the formation of immunothrombosis.

NE translocates into the nucleus and catalyzes chromatin decondensation, making it a pivotal component in the formation of NETs(Papayannopoulos et al. 2010). We observed that NE inhibition abrogates the development of HSOS in murine models, suggesting that NET is a key component of the pathophysiology of HSOS. Treatment with NE inhibitor or neutrophil depletion markedly attenuates neutrophil-platelet interactions, supporting that NETs mediate platelets sequestration in PA-HSOS. VWF released by damaged endothelial cells mediates NETs adhesion to liver sinusoids through a histone-VWF interaction. Simultaneously, NET-derived constituents inhibit ADAMTS13 activity, fostering the accumulation of ultra-large VWF multimers(Yang et al. 2020). In this study, NET inhibition correlates with reduced sinusoidal VWF signal intensity, confirming NETs’ role in promoting VWF fiber formation in PA-HSOS. The histone-DNA backbone of NETs is believed to add stability and rigidity to the fibrin scaffold in thrombin(Longstaff et al. 2013). Consistent with the mechanism, neutrophil depletion markedly attenuated fibrin deposition in HSOS mice, providing direct evidence that neutrophil is a critical amplifier of fibrin generation. Platelets can activate neutrophils to form NETs. Also, platelet could adhere to NETs and become activated, as NETs histone H3 and H4 could trigger platelet activation and aggregation in vitro(Wienkamp et al. 2022). In PA-HSOS livers this feedback loop is manifest as tight neutrophil-platelet co-clusters that dissolves upon neutrophils depletion or NETs inhibition, confirming the co-localization is NET-dependent. Collectively, our findings establish NETs as a central scaffold that orchestrates thrombus formation in PA-HSOS by promoting platelet recruitment, fibrin assembly, and VWF release, which provide a mechanistic explanation for hepatic platelet accumulation, fibrin deposition, and VWF up-regulation observed in PA-HSOS patients(Ji et al. 2024).

NETs can be broadly categorized as suicidal NETosis, in which neutrophils die after expelling the filaments; and viable NETs, in which the neutrophils remain vital and functionally competent after vesicles release. SytoxGreen, a cell-impermeant nucleic acid dye, is widely employed to quantify extracellular DNA and thereby serves as a surrogate indicator of NETs formation. The fluorescence signal in supernatant and cells could be used to discriminate cell-free (viable) versus anchored (suicidal) NETs(Tanaka et al. 2015). This study found that neither MCT nor its metabolite DHMC could directly induce the formation of cell-free NETs (no increase in cfDNA or MPO-DNA). However, soluble factors released by DHMC-injured HHSEC selectively promoted the formation of cell-free NETs rather than anchored ones, indicating that conditioned medium from DHMC-treated HHSEC potently elicits vital NETs formation. This effect was independent of platelets, suggesting that endothelial cell injury may play a key role in MCT-related NETs. Additionally, SEN metabolites generated by HLMs indirectly induced NETs via HHSEC-conditioned medium, accompanied by the formation of DHP-GSH adducts. The enzyme kinetic parameter further supported the role of metabolic activation in NETs. These findings suggest that PA toxicity may indirectly promote NETs through endothelial injury rather than direct neutrophil activation, highlighting a potential role of endothelial-derived mediators in NET-driven pathology.

Subsequently, we performed transcriptome sequencing to delineate the pathways governing NETs formation in SEN-HSOS. Compared with controls, widespread up-regulation of neutrophil chemoattractants was evident in livers of SEN-HSOS mice. Neutrophil directed recruitment, or so-called chemotaxis, is orchestrated by a series of guidance cues. CXCL1 to 3 and CXCL5 to 8 are the seven human neutrophil attracting and activating chemokines, with CXCL8 as the most potent protein(Cambier et al. 2023). CXCL8 is up-regulated in DHMC-treated HHSEC, indicating its involvement in neutrophil mobilization induced by DHPA. Rodents do not have the gene for CXCL8(Cambier et al. 2023), so we measure mRNA levels of the functional homologues and verified the corresponding protein expression(Griffith et al. 2014). Our results revealed that hepatic CXCL1 and CXCL2 were up-regulated in both PA-HSOS models. Notably, the expression of hepatic CXCL5 and CXCL7 was markedly elevated in rats yet remained unchanged in mice, underscoring a species-specific divergence in chemokine responses. CXCR2 is suggested to be a main mediator for neutrophil chemotaxis(Metzemaekers et al. 2020), and we observe the up-regulation of hepatic CXCR2, indicating CXCL1/2-CXCR2 interaction mediates neutrophil migration in PA-HSOS. Despite 78% amino acid identity with CXCR2, CXCR1 selectively triggers a phospholipase D (PLD)-dependent signaling pathway that drives neutrophil oxidative burst and subsequent NETs formation(Ha et al. 2017). Intriguingly, in this study, CXCR1 shows two bands: a slow migrating form (∼55 kDa) and a fast-migrating form (∼70 kDa), which are supposed to be phosphorylated forms. Phosphorylation initiates CXCR1 internalization, and provides a migration stop signal when the chemoattractant concentration is high(Rose et al. 2004). Reduction of NETs by targeting CXCR1/2 has been reported to reduce thrombosis and lung injury in experimental human and murine sepsis(Alsabani et al. 2022). So, we hypothesize that PA-stimulated LSECs release CXCL8 (CXCL1/2 in rodents), which engages CXCR1/2 on neutrophils, thereby initiating downstream signaling that drives NETs formation in HSOS.

## 5. Conclusion

Overall, our experiments propose a new mechanism of sinusoidal occlusion in PA-HSOS associated with locally recruited neutrophils forming NETs (Fig. 8). Preventing the recruitment of neutrophils or NET formation should reduce immunothrombosis without affecting hemostasis. Targeting NETs could be a promising therapeutic approach for PA-HSOS in clinic.

**Fig. 8.**
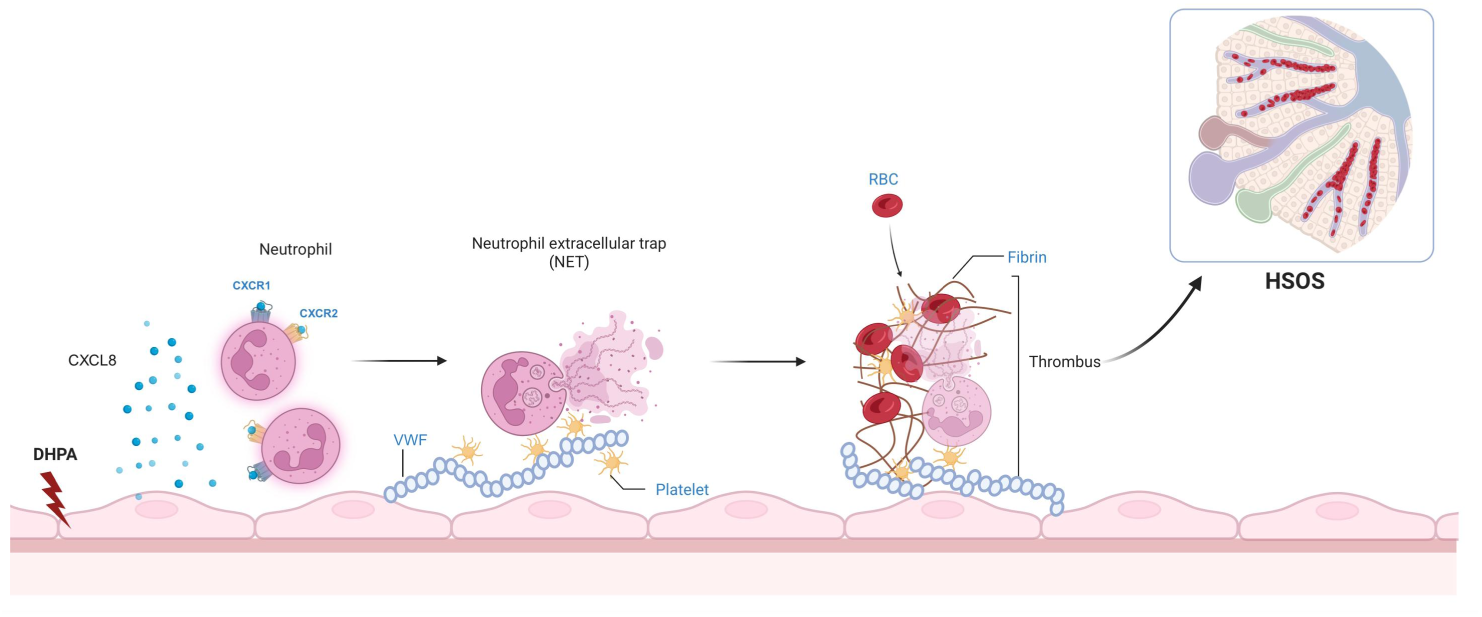
NETs participate in pathological development of PA-HSOS. Firstly, PA’s metabolite DHPA injures HHSECs, prompting them to release CXCL8 (CXCL1/2 in rodents). The resulting perisinusoidal CXCL gradient attracts neutrophils to the damaged endothelium via CXCR1/2. Activated neutrophils then generate NETs anchored to injured sinusoids by VWF. This in turn promotes VWF fiber formation, platelet accumulation, and further add stability to the fibrin scaffold in thrombi, thereby occluding the hepatic sinusoids. Image created with Biorender.com, with permission.

## Author contributions

Qiu Furong conceived the idea. Li Yue and Qiu Furong designed the study and provided the conceptual framework for the study. Zhang Shuang, Yan Dongming and Cheng Si performed the experiments, and analyzed the data. Cui Jiamin and Jin Jingyi participated in the animal experiments. Liu Chenghai provides resources. Yan Dongming and Li Yue wrote the manuscript. All authors commented on and approved the final version of the manuscript.

## Acknowledgements

This study was supported by the National Science Foundation of China [No.82574747, 82405066] and Opening Project of Shanghai Key Laboratory of Traditional Chinese Clinical Medicine (Shuguang Hospital) [No.20DZ2272200].

## Data availability

The data generated or analyzed during this study are included in this published article and its supplementary information files. The transcriptomic data are publicly available in Mendeley Data, doi: 10.17632/9xsxkmx9bs.1. Any other data sets that support the findings of this study are available on request from the corresponding author.

