## Supplementary Table. 1, Supplementary Fig. 1, Supplementary Fig. 2 for "Neutrophil extracellular traps-mediated thrombosis drive pyrrolizidine alkaloid-induced hepatic sinusoidal obstruction syndrome"

**Supplementary Data**

**Supplementary method**

**LC-MS/MS analysis**

GSH-DHP generated from HLMs system was detected by an LC-MS/MS system. The LC-MS/MS system contained an Agilent-1260 liquid chromatograph (Agilent Technologies, Santa Clara, CA) and an API 4000 triple quadrupole mass spectrometer (AB Sciex, Foster City, CA) equipped with an electrospray ionization source. The experiments were conducted with the electrospray source in positive ionization mode. The optimum parameters for the ionization source and the collision cell were capillary tension (4500 V), source temperature (500 ℃), nebulizer gas (50 psi), drying gas (50 psi), collision gas (medium), and curtain gas (10 psi). Chromatographic separation was performed on an Agilent Eclipse XDB-C18 reversed-phase column (150×4.6 mm, 5 μm) at 40 ℃. The mobile phase A was 0.1% formic acid and mobile phase B was acetonitrile containing 0.1% formic acid. The mobile phase was delivered at flow rate of 0.8 mL/min with gradient as follows: 0–1 min: 40% B; 1–3 min: 40%–80% B; 3–5 min: 80% B; 5–5.1 min: 80%–40% B; 5.1–10 min: 40% B. 2 μL of sample was injected to mass spectrometer to perform analysis using multiple reaction monitoring (MRM) of the transitions of m/z 336.0→m/z 94.0 for SEN, and m/z 425.0→m/z 118.0 for GSH-DHP, and m/z 392.2→m/z 264.3 for SHG, respectively. The metabolites were subjected to LS/MS analysis in neutral loss (NL) mode event. The NL ion of the m/z 307.3 was acquired with a collision energy of 20 to 50 eV, and the retention time of chromatographic peak was compared with those obtained from MRM mode.

**Supplementary figure**

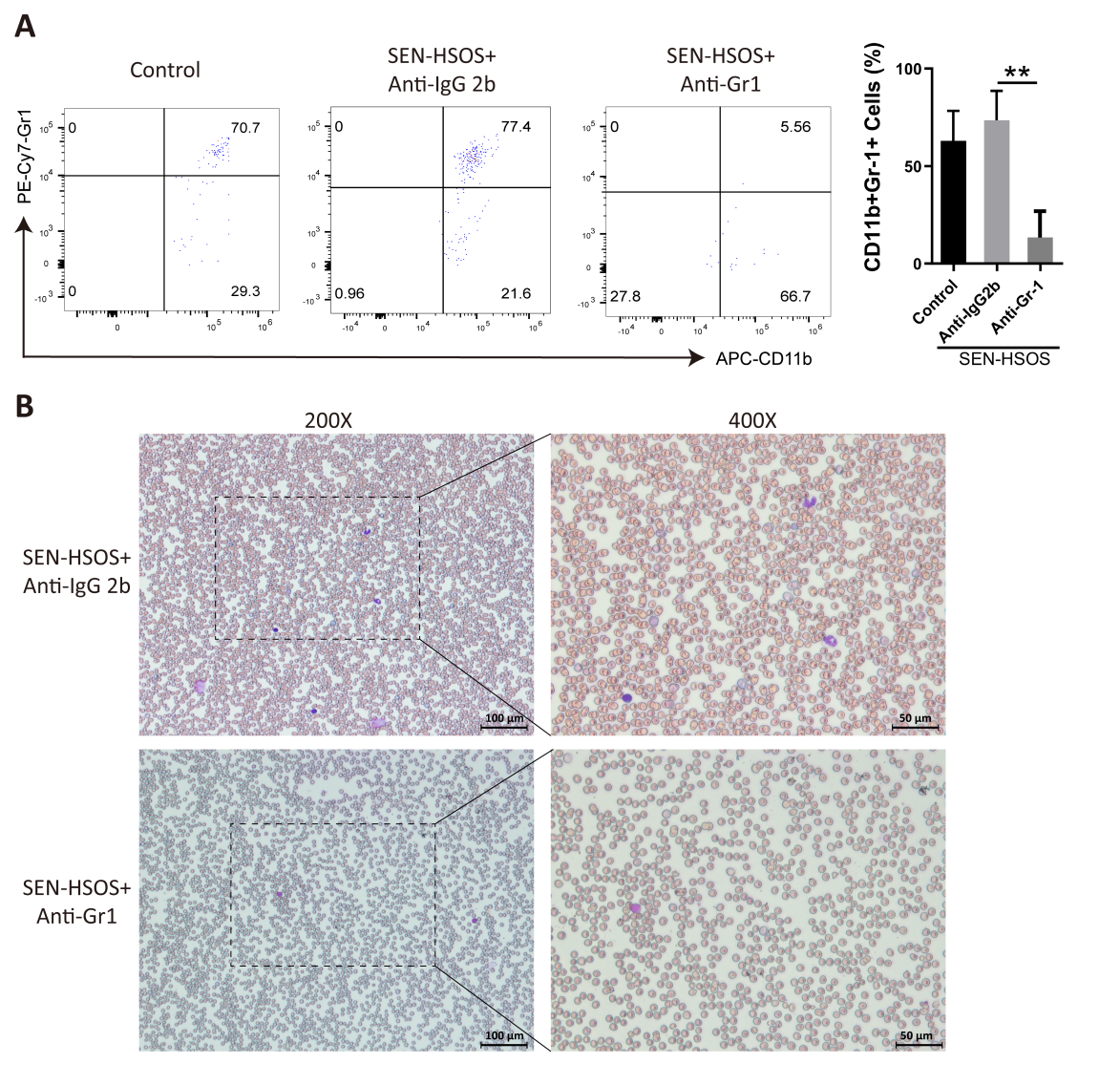

**Supplementary Figure 1** **Neutrophil depletion in the SEN-HSOS mouse model following anti-Gr1 antibody administration. (A)** Representative flow-cytometric dot plots illustrating the gating strategy for peripheral blood CD11b⁺Gr-1⁺ cells. Quantification (right panel) reveals a significant reduction in neutrophil frequency after anti-Gr1 treatment. **(B)** Representative Wright-Giemsa-stained peripheral blood smears corroborating neutrophil depletion (original magnification, ×200, ×400). Data are presented as mean±SD. ***p*< 0.05 vs control group, n = 3-6.

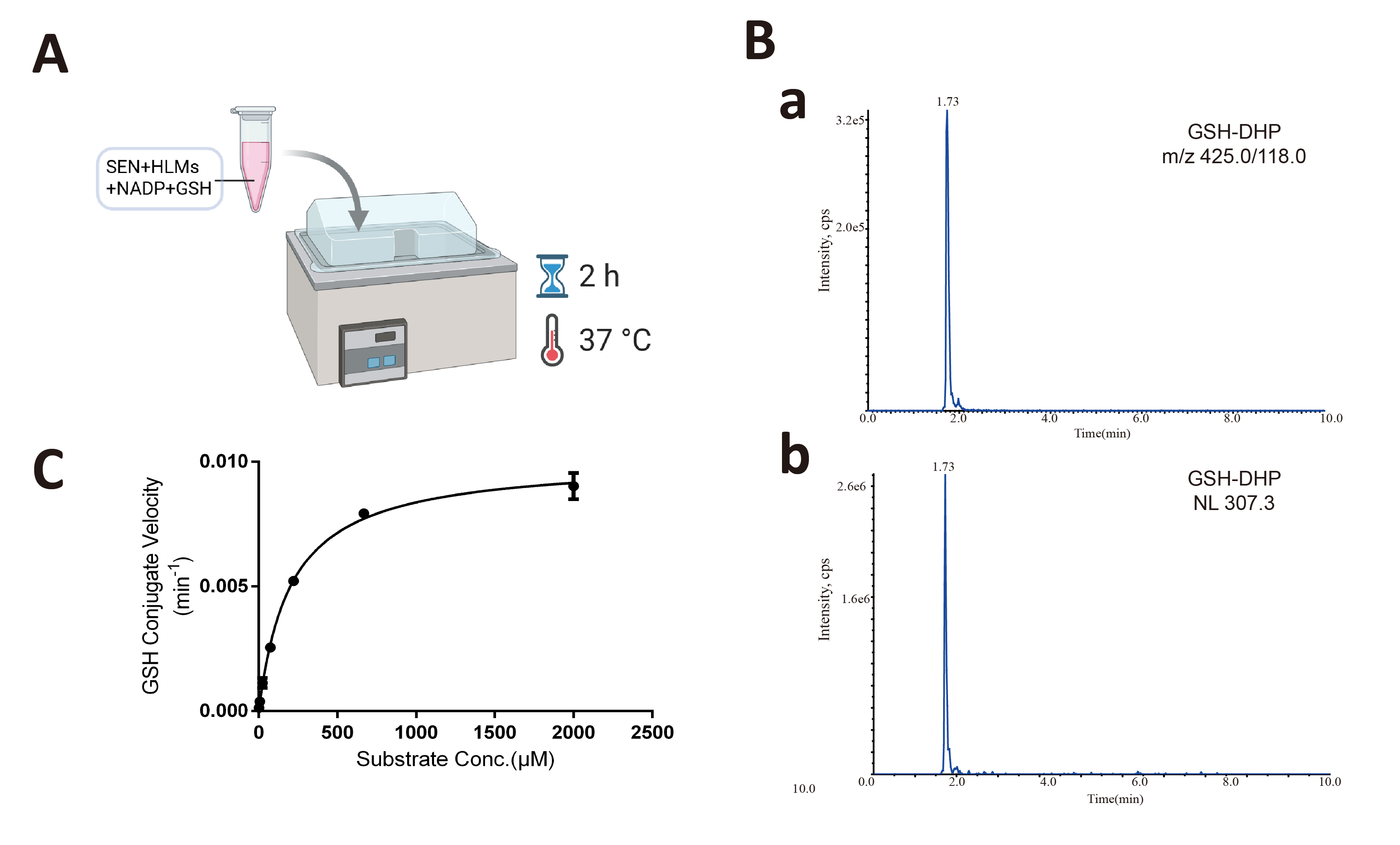

**Supplementary Figure 2 SEN metabolites generation in HLMs. (A)** Scheme of in vitro metabolic activation assay. **(B)** Representative LC-MS/MS chromatograms of the GSH-DHP conjugate: (a MRM transition m/z 425→118; b NL scan of 307.3). **(C)** Michaelis-Menten fitting curve of GSH-DHP formation.

**Supplementary table**

Supplementary Table1. Primers sets used for RT-PCR

| Gene | Primer sequences |
| --- | --- |
| CXCL1(KC) | Forward: GCTGGGATTCACCTCAAGAACATC |
|  | Reverse: GTGTGGCTATGACTTCGGTTTGG |
| CXCL2(MIP-2α) | Forward: CCACCAACCACCAGGCTACAG |
|  | Reverse: GGCTTCAGGGTCAAGGCAAAC |
| CXCL3(MIP-2β) | Forward: CACTGGTCCTGCTGCTGCTG |
|  | Reverse: CGTCACCGTCAAGCTCTGGATG |
| CXCL5(LIX) | Forward: CAGCTCCGCCCGCATCC |
|  | Reverse: GGCAGCGTGAACAGCAACAG |
| CXCL7(NAP-2) | Forward: AGACCTACATCGTCCTGCACCAG |
|  | Reverse: GGAGCCAGCGCAACAAGGATC |
